# Opto2P-FCM: A MEMS based miniature two-photon microscope with two-photon patterned optogenetic stimulation

**DOI:** 10.1101/2024.10.21.619528

**Authors:** Gregory L. Futia, Mo Zohrabi, Connor McCullough, Alec Teel, Fabio Simoes de Souza, Ryan Oroke, Eduardo J. Miscles, Baris N. Ozbay, Karl Kilborn, Victor M. Bright, Diego Restrepo, Juliet T. Gopinath, Emily A. Gibson

## Abstract

Multiphoton microscopy combined with optogenetic photostimulation is a powerful technique in neuroscience enabling precise control of cellular activity to determine the neural basis of behavior in a live animal. Two-photon patterned photostimulation has taken this further by allowing interrogation at the individual neuron level. However, it remains a challenge to implement imaging of neural activity with spatially patterned two-photon photostimulation in a freely moving animal. We developed a miniature microscope for high resolution two-photon fluorescence imaging with patterned two-photon optogenetic stimulation. The design incorporates a MEMS scanner for two-photon imaging and a second beam path for patterned two-photon excitation in a compact and lightweight design that can be head-attached to a freely moving animal. We demonstrate cell-specific optogenetics and high resolution MEMS based two-photon imaging in a freely moving mouse. The new capabilities of this miniature microscope design can enable cell-specific studies of behavior that can only be done in freely moving animals.

## Introduction

Miniaturized head-attached microscopes are valuable tools to measure the activation of neural networks *in vivo* within a behavioral context. These devices enable researchers to observe and analyze neural activity, providing insight into how neural circuits function in real time during various behaviors ^1–6^. Among these tools, multiphoton miniature microscopes offer greater depth access, higher resolution and eliminate out-of-focus background compared to miniature one-photon microscopes^7–14^. The enhanced imaging capabilities of miniature multiphoton microscopes make them a powerful tool for neuroscience research. Understanding neuronal activity mediating behavior is a fundamental question in neuroscience. This complex task requires not only advanced tools to record activity patterns but also precise manipulation of activity in specific neurons within the brain circuits of freely-behaving animals^15–19^. To study the behavioral relevance of neural activity, there is a growing interest in developing methods that integrate imaging and spatially patterned photostimulation into miniaturized microscopes.

To further this goal, we developed a miniature head-attached microscope using a MEMS based scanner for two-photon (2P) imaging combined with spatially patterned 2P photostimulation, termed the Opto2P-FCM (optogenetics two-photon fiber-coupled microscope). We demonstrate neuron level optogenetic stimulation and high resolution Ca^2+^ imaging in freely moving mice. The microscope design is illustrated in Fig. 1. The system includes two femtosecond pulsed lasers, at 920 nm and 1030 nm wavelengths, and a two-channel detection system. Excitation light at 920 nm is delivered to the head-attached microscope through a polarization maintaining (PM) fiber and imaging is performed with an on-board MEMS scanning mirror. The 1030 nm laser is spatially shaped using a spatial light modulator (SLM) and the pattern is propagated through a flexible coherent imaging fiber bundle (CFB) to a second beam path in the miniature microscope. The 1030 nm beam is combined with the 920 nm laser using a dichroic mirror and both are overlapped at the focus. Fluorescence from the sample is collected back through the objective, transmitted by the dichroic and focused onto the CFB, where it is relayed to the photodetectors. The miniature microscope design uses the same optical beam path for photostimulation and fluorescence collection which greatly reduces the complexity and weight. 2P imaging combined with 2P targeted photostimulation in the somatosensory and visual cortex in freely moving mice expressing jGCaMP7s and ChRmine^20^ is demonstrated (Fig. 1c). Photostimulation from selected regions in the field of view result in robust Ca^2+^ fluorescent transients timed to the stimulation pulse. These results are the first demonstration of a miniature head-mounted microscope that can perform high resolution 2P imaging using a MEMS scanner with 2P targeted photostimulation needed to record and manipulate specific neurons.

**Figure 1:**
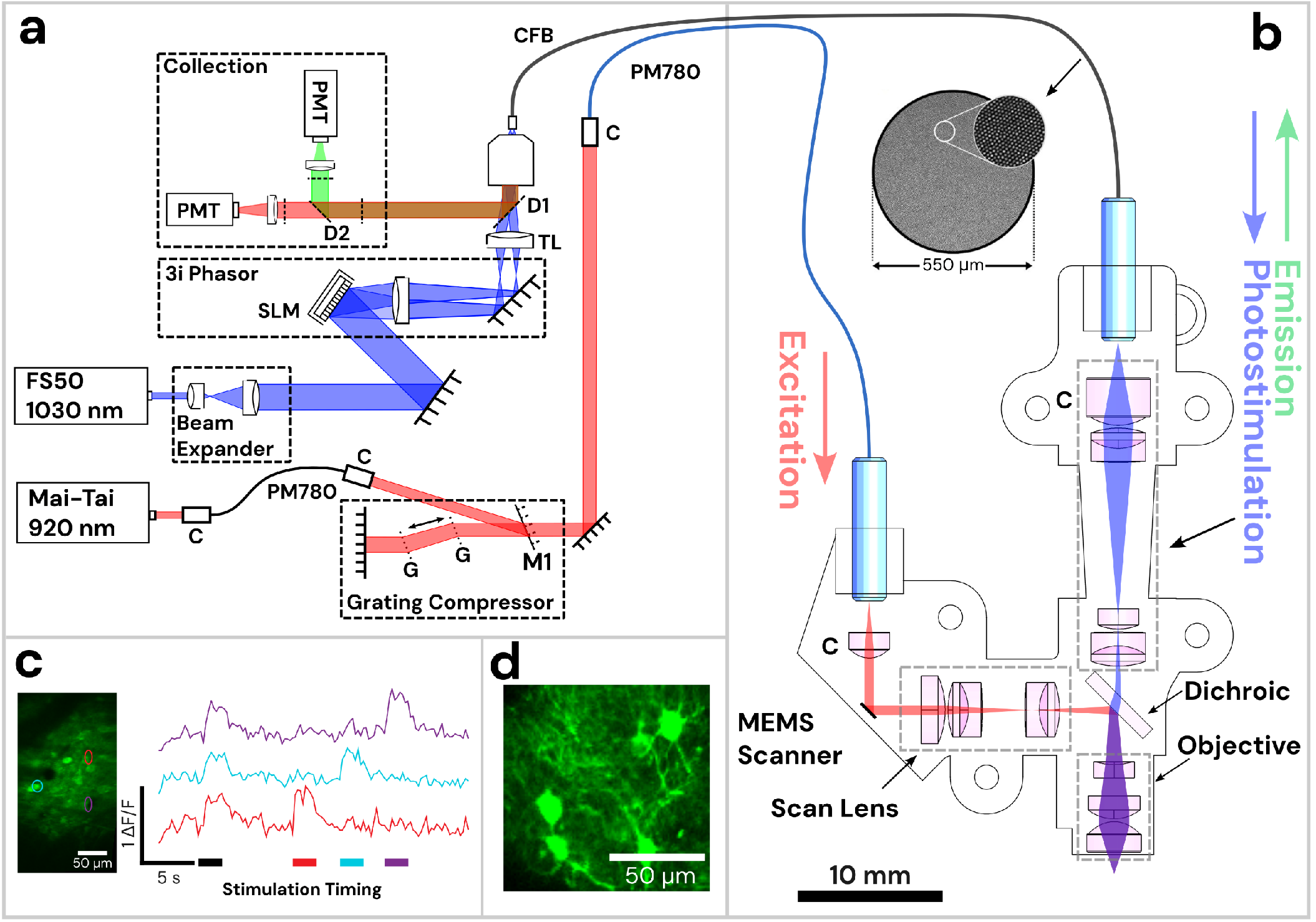
Diagram of Opto2P-FCM miniature two-photon microscope with optogenetic stimulation. (a) Schematic of the lasers and detection system. The 920 nm pulsed laser is coupled into a polarization maintaining fiber (PM780) for spectral broadening and a transmission grating-pair compressor for dispersion compensation. The output is coupled to a second PM780 fiber to relay to the miniature microscope. A 1030 nm pulsed laser is beam-expanded and reflected off of a spatial light modulator (SLM) for holographic spatial patterning. The spatially patterned beam is focused onto a coherent imaging fiber bundle (CFB) to relay the patterned light to the miniature microscope. Fluorescence is collected back through the same CFB, separated by dichroics and detected on two photomultipliers (PMTs). (b) Schematic of miniature microscope with detailed layout of optical components and beam path. The 920 nm laser output from the PMF is collimated and reflected off of a MEMS scanner and passes through a scan lens consisting of a plano-convex and achromatic lens to generate a telecentric scan across the imaging plane. Patterned photostimulation light at 1030 nm exits the CFB, is collimated using an aspheric lens, combined with the 920 nm light using a dichroic mirror and focused through the same objective. The miniature objective is designed with a working distance of 1.5 mm to focus both 920 nm and 1030 nm light to the same imaging plane. (c) In vivo demonstration of head-attached two-photon imaging and selective optogenetic two-photon photostimulation in a freely moving mouse. Imaging was performed in the visual cortex with neurons co-expressing the calcium indicator jGCaMP7s and the opsin ChRmine. Two-photon photostimulation was performed on three selected regions of interest (ROI) in the imaging field (see outlines on the maximum intensity projection image). An image time sequence was acquired whereby all three ROIs were illuminated simultaneously (indicated by black stimulation timing bar) followed by each individual ROI sequentially (indicated by color matched bars). Δ*F*/*F* traces of jGCaMP7s show responses in the ROIs that occur synchronously with photostimulation. (d) The resolution of the Opto2P-FCM provides visualization of processes from somatosensory cortex neurons expressing jGCaMP7s.

## Results

### Optical performance

The image resolution of the Opto2P-FCM was characterized using a thin fluorescent rhodamine sample spin-coated on a slide. The slide was scanned axially through the focus of the 920 nm excitation beam. Two-photon fluorescence was detected in transmission and the resulting trace is shown in Fig. 2a. The full-width-half-maximum (FWHM) was calculated to be 14 μm. The resolution of the 1030 nm photostimulation was measured in a similar manner. Two-photon fluorescence intensity as a function of axial position was measured for two different photostimulation patterns, 20 and 40 μm diameter circles, and found to have a FWHM of ∼50 and 54 μm respectively (shown in Fig. 2b and c). Additionally, the peak of the axial profiles for the 920 nm and 1030 nm beams overlapped with < 5 μm axial shift.

**Figure 2:**
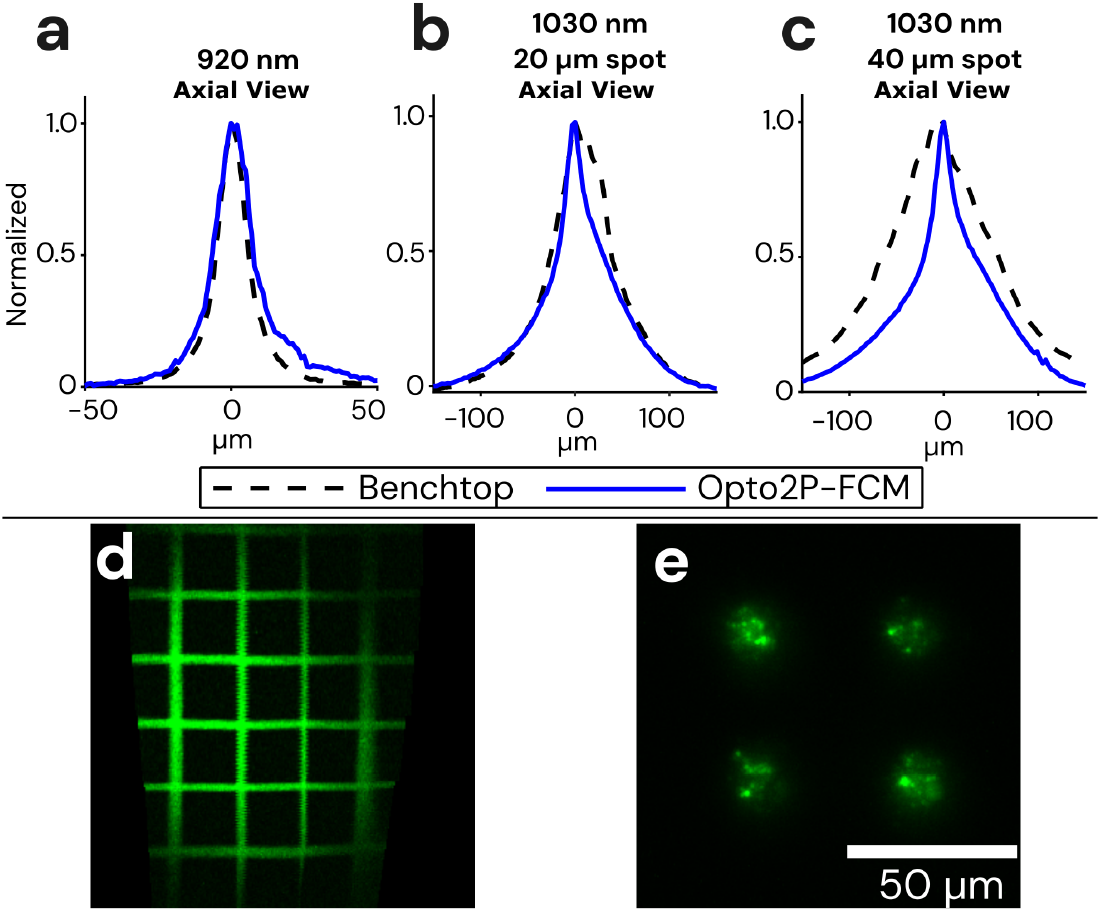
Optical resolution and field of view measurements of the Opto2P-FCM. (a) Axial resolution of the Opto2P-FCM was characterized by measuring the 2P fluorescence signal of a uniform flat fluorescent target scanned through the focus of the 920 nm laser (blue solid line, 14 μm FWHM) compared with a benchtop two-photon microscope with a 10x/0.4 NA objective (dashed line, 11 μm FWHM). Axial resolution of the 1030 nm patterned photostimulation was measured for a (b) 20 μm diameter circle using the Opto2P-FCM (blue solid line, 50 μm FWHM) compared to the benchtop microscope (dashed line, 54 μm FWHM) and for a (c) 40 μm diameter circle using the Opto2P-FCM (blue solid line, 54 μm FWHM) compared to the benchtop microscope (dashed line, 124 μm FWHM). (d) Opto2P-FCM image of a 50 μm grid slide showing a field of view of ∼200 μm × 300 μm. (e) Image of photostimulation pattern (4 circles with 20 μm diameter) at the focus of the Opto2P-FCM recorded by 2P fluorescence from a thin rhodamine sample.

For comparison, the resolution of an upright commercial two-photon microscope (Sutter Instruments Movable Objective Microscope) with a 10x/0.4 numerical aperture (NA) objective was measured in a similar manner. The fluorescence intensity axial profile on the benchtop system had a FWHM of 11 μm which is only slightly smaller than the Opto2P-FCM indicating a comparable numerical aperture. The axial resolution of the 1030 nm photostimulation beam was characterized for 20 and 40 μm diameter spots and measured to have a FWHM of 66 μm and 124 μm on the benchtop system. In comparison, the Opto2P-FCM has improved axial resolution with less dependence on lateral spot size. This effect is due propagation of the light through the multi-core CFB where light emitted from individual cores is no longer fully coherent^21–23^. The imaging field of view (FOV) was measured to be 200 μm × 300 μm by imaging a 50 μm grid slide calibration target (Max Levy II-VI Aerospace and Defense, DA113) (Fig. 2d).

### Fiber delivery

One benefit of the system is the use of standard polarization maintaining fiber (PM780-HP, Thorlabs) to deliver the short-pulsed 920 nm excitation light. In contrast, other miniature 2P and 3P microscopes use hollow core fibers which are significantly more costly and require expertise to connectorize. In order to obtain pulses with high peak powers at the output of the fiber, the 920 nm laser is first spectrally broadened using a 15 cm length PM fiber, sent to a transmission grating-pair compressor that applies 100,000 fs^2^ of anomalous dispersion and coupled into a second PM fiber to deliver the 920 nm light to the miniature microscope. The addition of the 15 cm PM fiber avoids spectral narrowing assocated with nonlinear self-phase modulation^24,25^ allowing a short pulse at the end of the fiber, as demonstrated in our prior publication^7^. The 920 nm output from the fiber was characterized using frequency resolved optical gating (FROGscan, MesaPhotonics) with a measured pulse of 134 fs FWHM and 8 nm spectral bandwidth (see Supplementary Fig. S4). The average power used for *in vivo* 2P imaging ranged from 15-30 mW with associated peak powers of 1.5-3.7 kW.

### Optogenetic stimulation and recording in freely moving mice

We demonstrate simultaneous MEMS based 2P calcium imaging and 2P patterned optogenetic stimulation in fiber-tethered freely moving mice with the Opto2P-FCM (Fig. 3). C57BL/6 mice were injected with AAVs for expression of the calcium reporter jGCaMP7s and opsin ChRmine and implanted with 4 × 4 mm^2^ cranial windows^26^ positioned over the visual cortex or somatosensory cortex. A 3D printed baseplate was attached to the animal’s head with dental cement. The Opto2P-FCM was held to the baseplate during the experiment. Helium filled balloons were attached with a fishing line to a tie off point on the device housing to provide a counterweight, yielding an effective weight of ∼3 grams. Mice were then placed in a cage and allowed to move freely and behavioral video was captured by an IR illuminated camera. The mice were able to navigate the full extent of the cage and freely engage with a running wheel (Fig. 3c and f and Video S1). Multi-region patterned optogenetic stimulation was performed on 6 regions of interest (ROIs) with different timing patterns that included stimulation of just one ROI at a time and stimulation of all ROIs simultaneously. The fluorescence signal from jGCaMP7s was recorded simultaneously. Imaging data was analyzed using CaImAn^24^ to identify regions of jGCaMP7s transients (Fig. 3a and d) and extract Δ*F*/*F* traces (Fig. 3b and e). These results showed good overlap with the stimulation ROIs and timing of fluorescence transients. Denoised timelapses from CaImAn and raw images are shown in Video S1. Robust responses to individual ROI stimulations were consistently observed, while stimulations of all ROIs simultaneously resulted in varying response between cells. The variability in multi-region stimulation responses can be attributed to different expression levels of calcium indicator and opsin and core-to-core inhomogeneity of two-photon excitation efficiency in the CFB^27–29,21,22^. More consistent stimulation should be possible by weighting the intensities between the different ROIs. The experiments were performed over multiple days with the ability to return to the same imaging field (Fig. 4).

**Figure 3:**
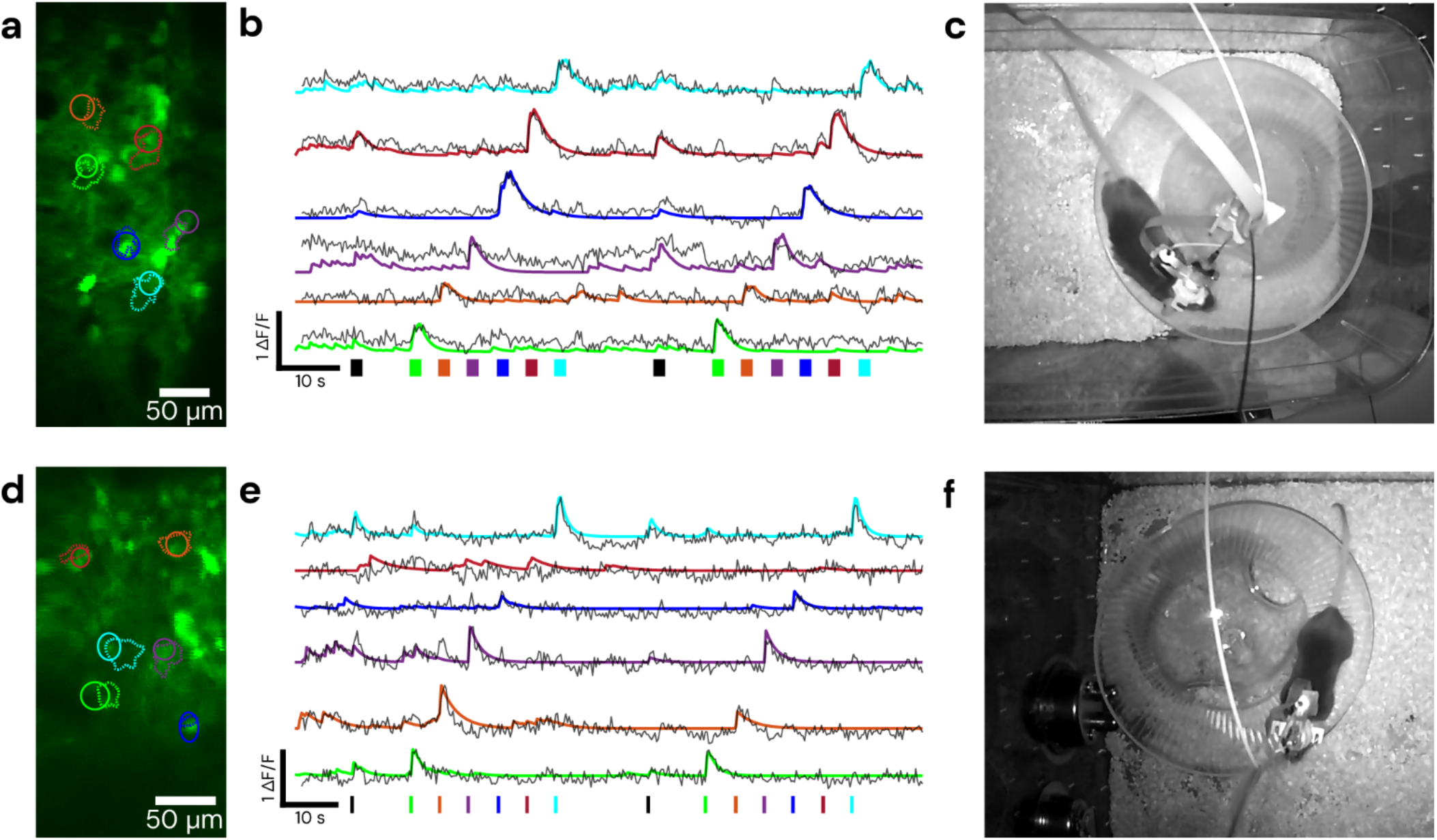
Demonstration of cell-specific 2P patterned photostimulation and simultaneous 2P imaging in a freely moving mouse. The stimulation sequence consists of all ROIs together followed by stimulation of each ROI individually. Top panels were taken using a 0.5 m length CFB in visual cortex, with 5 ms/frame of stimulation for 2 seconds, bottom panels were taken using a 1 m length CFB in somatosensory cortex, with 10 ms/frame of stimulation for 700 ms. (a) and (d) show the fields and targeted/detected ROIs, with solid line circular ROIs indicating the targeted stimulation regions and dashed line ROIs indicating those detected by CaImAn. (b) and (e) show Δ*F*/*F* traces of jGCaMP7s activity during the recording timelapse color matched to ROIs in (a) and (d). Solid lines represent Δ*F*/*F* denoised traces from CaImAn and overlaid dashed lines are raw Δ*F*/*F* traces of the stimulation ROI. Color matched bars at the bottom represent stimulation times and durations, with black representing the stimulation of all ROIs. (c) and (f) show a still frame of the mouse during the recording. Behavioral video is shown in Video S1.

**Figure 4:**
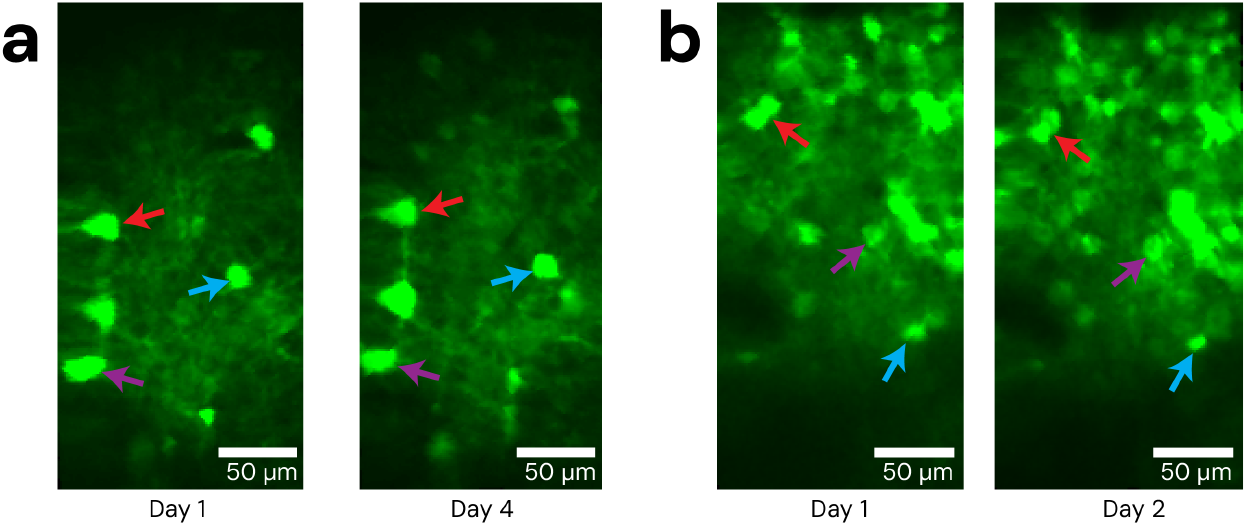
Repeatability of Opto2P-FCM imaging fields. The Opto2P-FCM was head-attached to moving animals across multiple days, repeatably accessing the same cell field after being locked into the baseplate. (a) Example of repeated imaging of the same field across days in visual cortex and (b) somatosensory cortex. Colored arrows indicate several reference features.

## Discussion

In this study, we accomplished simultaneous two-photon imaging and two-photon patterned photostimulation of neural activity in visual and somatosensory cortex of freely behaving animals using Opto2P-FCM, a MEMS based miniature microscope. We showed repeated imaging of the same cell field over multiple days. The ability to simultaneously image and photostimulate neural activity in neuronal soma of freely moving animals using this MEMS scanner-based device represents a significant advancement over previous miniature microscope designs. The Opto2P-FCM can resolve neuronal features such as dendritic structures as shown in Fig 1d. The novel design of the Opto2P-FCM optics combines the photostimulation and fluorescence collection pathways making the device lighter, more compact and requiring fewer optics. An additional innovation is the use of pulse compression techniques to overcome nonlinear self-phase modulation to allow use of a standard glass fiber. We demonstrate short femtosecond pulses after the standard PM fiber for efficient 2P excitation. In addition to being much lower cost and commercially available, PM fibers have reduced stiffness and are less brittle compared to the hollow core fibers used in other multiphoton miniature microscopes^10,12,14^. The PM fiber also allows the Opto2P-FCM to be adapted to different excitation wavelengths, which are not supported in hollow core fibers that are designed for a narrow range of wavelengths. Additionally, collection of the fluorescence emission through the fiber bundle allows multiple color channels to be detected simultaneously without adding the weight of a second photodetector on the head attachment.

The Opto2P-FCM has benefits over other miniature microscopes and microendoscopes that have been developed for combined imaging and spatially patterned photostimulation. A one-photon endoscope was demonstrated using a miniature digital micromirror device (DMD) for spatial patterning of the photo-stimulation^30^. However, the integration of a DMD adds additional weight (∼7.8 g) and one-photon photostimulation activates neurons above and below the focal plane. Another demonstration used a CFB coupled to a gradient index of refraction (GRIN) microendoscope lens to demonstrate freely moving 2P imaging and photostimulation^23^. However, 2P imaging through CFBs are intrinsically limited in their resolution due to core-to-core pixelation making it impossible to image cellular processes. Additionally, CFBs are well known to have core-to-core variation in 2P excitation efficiency and polarization which degrades 2P fluorescence imaging through CFBs ^22^. The Opto2P-FCM with a MEMS scanner overcomes these limitations by fully utilizing the resolution of the objective.

We anticipate that the Opto2P-FCM will provide a novel neuroscience tool that can advance our ability to conduct detailed mechanistic investigations of neural activity in freely behaving animals. By allowing researchers to observe and manipulate neural circuits in real-time, while an animal engages in natural behaviors, this technology will provide deeper insights into the neural basis of behavior. The Opto2P-FCM could lead to a better understanding of how specific neural pathways contribute to cognitive processes, sensory perception, motor control, emotion^31^, cognition^32,33^ and learning^34^, ultimately opening new avenues for research in neuroscience and related fields. An example of a future application of this device for a behavior requiring freely moving animal interactions is the study of the neural basis underlying social bonding^35–37^ and cognition. How these social bonds are represented in the brain is a question that has been previously tackled using cellular resolution Ca^2+^ imaging^38^. To move from descriptive to functional approaches, a method is needed to specifically activate or inhibit subsets of neurons in the context of behavior with prospective mates. Another application is the study of place cells in the hippocampus that encode for spatial information during navigation and are key to understand the neural basis of learning and memory^39^. Major advances in understanding place cells has been done by imaging their activity through two photon imaging in head fixed animals navigating a scene in virtual reality (VR)^40^, but two dimensional place tuning was found to be profoundly impaired in VR navigation ^41^ and the theta rhythm frequency has been found to be slower in VR environments^42^. The role of place cells in spatial navigation has been studied in freely moving mice with one photon and two-photon miniature microscopes^10,43^, but did not include capabilities for optogenetic modulation. Combining imaging with optogenetics can provide an understanding the involvement of cellular processes in signal processing in place cells^44^.

Opto2P-FCM can enable optogenetic modulation of specific subsets of neurons or specific dendritic processes and simultaneous imaging of circuit activity at timed to behavioral epochs, allowing investigators to ask how activation or inhibition of cells or cellular processes affects behaviors such as social bonding and spatial navigation. One such application would be in the investigation of CA1 pyramidal neurons in spatial and epoch behavior. A recent study performed targeted holographic stimulation in subsets of cells in CA1 in head-fixed mice undergoing one dimensional spatial navigation to show that place cells drive memory-guided spatial behavior^45^. However, it remains a question if the activity of small subsets of neurons is causal in 2D spatial navigation. Using Opto2P-FCM, it would be possible to identify cells whose activity can be used to decode animal decisions in 2D spatial navigation tasks and stimulate these cells in a manner recreating the naturally observed activity. The Opto2P-FCM further advances the development of optical tools for neuroscience studies in freely behaving animals by allowing high resolution MEMS 2P imaging with cell-specific patterned 2P photostimulation for the first time. Miniature microscope designs have opened new possibilities for neuroscience studies no longer confined to head fixation under large benchtop microscopes. Developments of these miniature microscopes are rapidly progressing and the Opto2P-FCM adds new capabilities to these systems.

## Methods

### Optical design

The optical design for the Opto2P-FCM was carried out using ray tracing software, Zemax (Optics Studio), constrained to commercially available optics. The optical configuration of the 920 nm excitation beam path is similar to those of previously developed multiphoton microscopes^10^ with the exception of the use of only commercially available lenses. Novel to our design is the incorporation of holographic patterned 2P photostimulation and fluorescence collection path. The optical design and ray schematic are shown in Fig S1a. A clamshell housing was designed in SolidWorks to simplify the installation of the optics. The housing was printed using stereolithography (Formlabs Form 3+, Tough 1500 resin) as shown in Fig. S1b.

The 920 nm excitation light is delivered through a single mode PM fiber (PM780-HP, Thorlabs) and collimated using an aspheric lens and reflected from a MEMS scanner (Mirrorcle A3I12.2-1200AL) mounted on a custom designed PCB with flex cable. A scan lens comprised of plano-convex and achromatic lenses generates a telecentric scan as shown in the ray diagram, Fig. S1a, for scan angles of -4, 0 and 4 degrees followed by a dichroic and an imaging objective lens. The Opto2P-FCM is designed with an NA of 0.4 for 920 and 1030 nm with a magnification of ∼5.5X and 2.8X respectively and overlapping FOV of 250 μm × 250 μm. The design has a working distance of ∼1.5 mm (see Supplementary Fig. S1c) while maintaining a Strehl ratio of ∼0.8 across the scanning angles of -4 to 4 degrees. The point-spread function for scanning angles of 0, 2.5 and 5 degrees are shown in Fig. S2a with Strehl ratio of 0.8 and the focal spot within the Airy disk radius of 1.38 μm (dashed circle) for all scanning angles, demonstrating diffraction-limited performance. A custom dichroic (T ≥ 90% 330-870, 975-1200 nm – R ≥ 90% 895-935nm) separates the excitation and photostimulation/emission beam paths. The photostimulation/emission path is configured to be chromatically corrected and focuses the 1030 nm photostimulation light from the CFB (Fujikura FIGH-15-600N) to the same imaging plane as the 920 nm light. The point-spread function (PSF) of the 1030 nm light at the imaging plane is shown in Fig. S2b for an input beam launched at -200, 0, and 200 μm at the entrance facet of the CFB with 0.2 NA. The dashed circle with a radius of 1.73 μm shows the Airy disk. The Strehl ratio of the 1030 nm photostimulation beam varies across the imaging field with a range of 0.6 to 0.7. The root-mean square (RMS) radius of the spot is plotted for imaging (920 nm) and photostimulation (1030 nm) as a function of working distance with zero position on the minimum RMS spot at 920 nm (Fig. S2c). The minima of the RMS spots are offset by less than ∼0.5 μm. The density plot of the Strehl ratio for the excitation beam is shown in Fig. S2d as a function of scanning angle of the MEMS scanner and the working distance. The system shows a 2 μm axial displacement between the 920 and 1030 nm light when scanning to 4-degree angle relative to on-axis position.

### Inspection scope

An inspection scope was used to measure the axial resolution of the excitation and photostimulation beams, perform spatial calibration of the photostimulation and imaging fields, and ensure that the fields are parfocal. The inspection scope consists of a 10x/.4 NA Olympus objective, a 90 deg folding mirror, and a 100 mm focal length lens and camera (FLIR CM3-US-31S4M-CS, Edmund Optics). For imaging fluorescence emission a series of filters (BG39, short pass 745, band pass 617/73) were placed before the camera to reject the fundamental lasers at 920 and 1030 nm. For imaging the 920 nm and 1030 nm beams directly, OD filters were placed before the camera. Further details are provided in Fig. S3.

### Opto2P-FCM assembly

The optical configuration and the assembled Opto2P-FCM are shown in Fig. 1b and Fig. S1b. The headpiece weighs ∼5 grams with an overall dimension of 40 × 22 × 12 mm^3^. The optical elements are press-fitted into one half of the housing shell, while the other half is secured in place using 0-80 screws. The 920 nm excitation light is delivered by a polarization maintaining fiber (PM780) with a 2.5 mm bare ceramic ferrule on one end and an FC/APC connector on the other end. The light emerging from the PM fiber is collimated by an aspheric lens. The PM fiber is axially adjusted to ensure the outgoing beam is collimated by observing the output at both near and far distances (∼10 cm and ∼2 meters). After mounting the MEMS mirror, its position is fine-tuned by observing the reflected beam at far distances and then glued in the housing.

The imaging and photostimulation planes were set to be parfocal though adjustment of the axial position of the CFB in the housing. This was performed by mounting the device to a micromanipulator (MP-285, Sutter Instruments) for precisely controlling the axial scanning. The CFB was mounted to a manual translation stage for separate adjustment. A thin fluorescent sample was prepared by spin coating rhodamine onto a microscope slide. The slide was placed on a stage at the focus of the inspection scope. The position of the Opto2P-FCM focus for the 920 nm beam was measured by recording the 2P fluorescence on the camera at different axial positions using the Sutter stage and locating the position of maximum signal. The CFB was then axially adjusted with the manual translation stage to bring the focus of the 1030 nm light to the same position. The CFB was then fixed into place in the microscope housing using glue.

### Distortion correction and spatial calibration

The MEM scanner in the Opto2P-FCM is subject to scan field distortions resulting in distortions in the image. These distortions are caused by the fact that the two scan axes are coupled^46^. As one axis is actuated, crosstalk into the other axis occurs. This crosstalk results in y position deviation as the fast x-axis is actuated. We utilized software developed by Zong *et al*.^10^ to compute a correction matrix by first imaging a grid slide and selecting a set of anchor points. The software then computes a 2D piecewise linear transformation which is saved as a correction matrix. This correction matrix is imported into SlideBook imaging software (SlideBook 2024, Intelligent Imaging Innovations, Inc.) to compute a voltage-to-pixel mapping for real-time data acquisition. To perform spatial calibration of the photostimulation patterning, the Opto2P-FCM was mounted to the Sutter micromanipulator and the inspection scope was used. A grid slide was set on a sample holder and brought into the axial focus of the inspection scope. The grid slide was translated to one corner of the grid for unique identification. The same corner was then located by imaging with the Opto2P-FCM and the Sutter stage was translated to put the corner in the top left of the imaging field. The photostimulation spatial calibration function in SlideBook software was used to illuminate three calibration spots on different locations of the grid slide, imaged with the inspection scope. These spots were then identified in the imaging field of the Opto2P-FCM. Once located, the software computes an affine transformation between the holographic coordinate system and the geometrically corrected imaging coordinate system. The calibration was tested by placing targeted holographic spots on known positions of the grid slide and observing their location on the inspection scope.

### Optical setup

A diagram of the optical setup is shown in Figure 1a. Imaging beam path: The 920 nm 80 MHz repetition rate laser (MaiTai HP DeepSee) passes through an isolator (714, Conoptics) to prevent back reflection to the laser. The output is coupled into a 15 cm PM fiber to broaden its spectrum (Thorlabs PM780-HP) with an adjustable fiber collimator (Thorlabs ZC618APC-B) for maximum throughput. The broadening fiber output was collimated through a second fiber collimator and sent into a Treacy grating pair pulse compressor built from two high efficiency 600 line per mm transmission gratings (Wasatch Photonics) to compensate for up to ∼100,000 fs^2^ dispersion. The beam is coupled into a 2 m long PM fiber (Thorlabs PM780-HP) to deliver the excitation light to the Opto2P-FCM. Photostimulation beam path: The 1030 nm photostimulation laser (NKT Photonics, aeroPULSE FS50) goes through a beam expander (*f*_*1*_=20 mm, *f*_*2*_=75mm), and is then incident on a spatial light modulator (SLM) (X10468, Hamamatsu) within the Phasor module (Intelligent Imaging Innovations, Inc.). After the SLM, the beam is focused with a 200 mm lens in a 2f geometry, forming the hologram pattern. The pattern is then relayed with two 4f telescopes forming a conjugate image at the focal plane of a tube lens in a Olympus IX71 microscope chassis. The tube lens and objective (10x/0.35 NA) relays the image to the focal plane of the objective where it is focused onto the facet of the CFB. The CFB delivers the patterned light to the miniature microscope. The 1030 nm laser was operated at a 2 MHz repetition rate. Detection Path: The fluorescence emission relayed from the miniature microscope through the CFB, is collimated by the objective. The collimated beam reflects off of a dichroic mirror (LP 850), passes through a short pass filter (BG 39, Thorlabs), dichroic (D2) (FF552-DI02, Semrock) and band pass filters (510/42, 617/73) and focused on two photomultipliers (PMTs) (Hamamatsu H7422P-40) for simultaneous two color imaging (Fig. S5).

### Data acquisition and stimulation parameters

SlideBook 2024 (Intelligent Imaging Innovations, Inc.) software was used to control the MEMS scanning mirror, acquire images, calibrate and compute the spatial hologram patterns, and control the timing of photo-stimulation. The x and y galvo output channels were connected to the analog input board (BDQ PicoAmp 5.4, Mirrorcle Technologies Inc.) which connects to the MEMS mirror. The acquisition frame rate was 3.4 Hz. The photostimulation duration and power are controlled by an analog output from Slidebook connected to the power modulation input on the 1030 nm laser. The pulse timing signal of the photo-stimulation is controlled by a function generator triggered by the frame trigger output from SlideBook to provide one pulse per frame. The pulse timing signal goes to the pulse enable line on the laser. Pulse durations of 5 and 10 ms corresponding to duty cycles of 1.7% and 3.4%, were used in the experiments. The same timing signal was also used to gate the PMTs during photo-stimulation to prevent overload from background signal. The stimulation power measured at the focus of the Opto2P-FCM for individual ROIs was between 2.7 and 7.8 mW when modulated, which corresponds to 153 mW to 229 mW unmodulated.

### Animals

C57BL/6J (JAX RRID:IMSR_JAX:000664) and T29-1 (JAX RRID:IMSR_JAX:005359) mice were housed in a vivarium with a reversed light cycle of 14/10 hr light/dark periods with lights on at 6 a.m. Food (Teklad Global Rodent Diet No. 2918; Harlan) was available *ad libitum*. All experiments were performed according to protocols approved by the University of Colorado Anschutz Medical Campus Institutional Animal Care and Use Committee.

### Surgery

Mice 2 to 5 months of age were maintained under anesthesia by exposure to isoflurane (2-5%) and placed in a stereotaxic frame. A 4 × 4 mm square craniotomy was made above the target site using a dental drill. 650 nL of a virus cocktail consisting of a 2:1 ratio of pGP-AAV9-syn-jGCaMP7s-WPRE and pAAV8-hSyn-ChRmine-mScarlet-Kv2.1-WPRE, both diluted 10x from stock to 1×10^12^ vg/mL, was injected at the target site at a rate of 100 nL per minute. Target sites were in somatosensory cortex (From bregma: -1.5 mm posterior, 1 mm lateral, 0.3 mm ventral) or visual cortex (From bregma: –3 mm posterior, 2 mm lateral, 0.3 mm ventral). After the viral injection, a hand cut 4 × 4 mm cranial window made of precision #1.5 cover glass (Thorlabs CG15KH1) was placed to cover the exposed brain tissue. The perimeter of the cranial window was then sealed with 3M Vetbond tissue adhesive (3M ID B00016067). A head-fixing bar was placed over the cranial window, and C&B Metabond (Parkell SKU S380) was used to cement the head-fixing bar in place and further seal the cranial window. Mice were allowed to recover for 2 weeks before the initiation of experiments and to allow time for virus expression.

### Locating cell fields

Benchtop multiphoton microscopes typically use a camera for widefield fluorescence imaging to coarsely position and locate the imaging fields. To locate target fields with the Opto2P-FCM, animals were first head-fixed and imaged under an upright two-photon Sutter Instruments Movable Objective Microscope (MOM). A fluorescent fiducial marker was placed on the Metabond outside of the cranial window with a yellow Sharpie fine-tipped marker (Sharpie Item # 75847), which is then found with two-photon excitation at 920 nm. Target location positions were recorded relative to the fiducial marker using the Sutter micromanipulator coordinates. The head-fixed animal was then transitioned to the Opto2P-FCM recording setup, where the Opto2P-FCM was held by a Sutter Instruments micromanipulator (Sutter MP-285). The fiducial was located with the Opto2P-FCM and was then translated to the previously recorded target region coordinates for experiments.

### Head attachment

The Opto2P-FCM is compatible with base plates designed by Zong *et al*.^10^. For head mounting, the baseplate was first attached to the Opto2P-FCM and mounted to a Sutter micromanipulator. The field of interest was located in the head-fixed animal using the method described above. The baseplate was then secured in place on the animal’s head using Metabond dental cement. Setscrews (0-80) were used to lock the Opto2P-FCM into position in the baseplate. After the experiments, the Opto2P-FCM is detached from the baseplate and a 3D printed cover is locked into the baseplate to minimize debris on the cranial window. When placing the Opto2P-FCM in an already prepared baseplate, the animal is first head-fixed, then the Opto2P-FCM is inserted into the baseplate, the baseplate setscrews are tightened to lock the device in place, and then the animal is removed from the head-fixing apparatus.

### Freely moving recording

To perform freely moving experiments, two helium filled balloons are attached with fishing line to a tie off point on the Opto2P-FCM. The animal is removed from the head-fixing apparatus and placed in a 7.25” × 11.5” cage. Behavioral video is captured using a camera with built in IR illumination. Freely moving recording sessions lasted ∼1 hour. After the freely moving recording session, the animal is head-fixed again to loosen setscrews and remove the Opto2P-FCM.

## Data processing

Imaging data was processed for motion correction and calcium signal extraction using CaImAn^47^.

## Acknowledgements

The authors would like to thank the following individuals for assistance and insightful discussions: Professor Robert McLeod, Dr. Omkar Supekar, Diane Jung and Samuel Gilinsky for assistance with custom PCB modeling of the MEMS mirror. The authors would like to thank NKT Photonics for providing a demo of the aeroPULSE FS50 laser used to collect the data.

## Author contributions

G.F., M.Z., and C.M. performed the experiments, analyzed data and generated the figures. M.Z. designed, fabricated, and assembled the miniature head-attached Opto2P-FCM. G.F. designed and assembled the optical and detection system and performed characterization of the system. C.M. did baseplate fabrication and alignment and led the *in vivo* testing. F.S. and A.T. performed surgeries and prepared animals. R.O and E.M. performed mechanical design of the housing. K.K. and B.O. wrote data acquisition software. G.F., M.Z., C.M., V.M., K.K., B.O., D.R., J.G., and E.G. contributed to manuscript writing. All authors participated in discussions and data interpretation.

