## Supplemental Tables and Figures for "Opto2P-FCM: A MEMS based miniature two-photon microscope with two-photon patterned optogenetic stimulation"

**Table 1: Opto2P-FCM components**

Part numbers correspond to the labels in the schematic in Figure S1.

| Part number | Item number | Description | Supplier |
|---|---|---|---|
| 1 | 16684 | Asphere - Collimator | Edmund optics |
| 2 | 47859 | Plano-convex – Scan lens | Edmund optics |
| 3 | L-MAC001 | Achromat – Scan lens | Ross optical |
| 4 | L-MAC001 | Achromat | Ross optical |
| 5 | 83710 | Asphere - Collimator | Edmund optics |
| 6 | L-MAC001 | Achromat – Relay lens | Ross optical |
| 7 | 49177 | Plano convex – Relay lens | Edmund optics |
| 8 | L-MAC001 | Achromat – Relay lens | Ross optical |
| 9 | ZT915dcrb | Dichroic –<br>T ≥ 90% 330-870, 975-1200 nm<br>R ≥ 90% 895-935nm | Chroma technology |
| 10 | 84380 | Plano concave - Objective | Edmund optics |
| 11 | L-MPX009 | Plano convex – Objective | Ross optical |
| 12 | 83627 | Asphere – Objective | Edmund optics |
| MEMS scanner | A3I12.2-1200AL | Scanning mirror | Mirrorcle |
| Excitation fiber | PM780 | Deliver 920 nm light | Thorlabs |
| Coherent fiber bundle | FIGH-15-600N | Deliver 1030 nm light and collect the emitted fluorescence light | Fujikura |

### Table 2: Laser Specifications

| Part | Description | Wavelength (nm) | Pulse Duration (fs) | Repetition rate (MHz) | Max Output Power (W) |
|---|---|---|---|---|---|
| SpectraPhysics Mai Tai HP DeepSee | Ultrafast laser for two-photon laser scanning microscopy | 920 | 80 | 80 | 1.85 |
| NKT Photonics aeroPULSE SF50 | Ultrafast laser for two-photon photostimulation | 1030 | 400 | 2 | 50 |

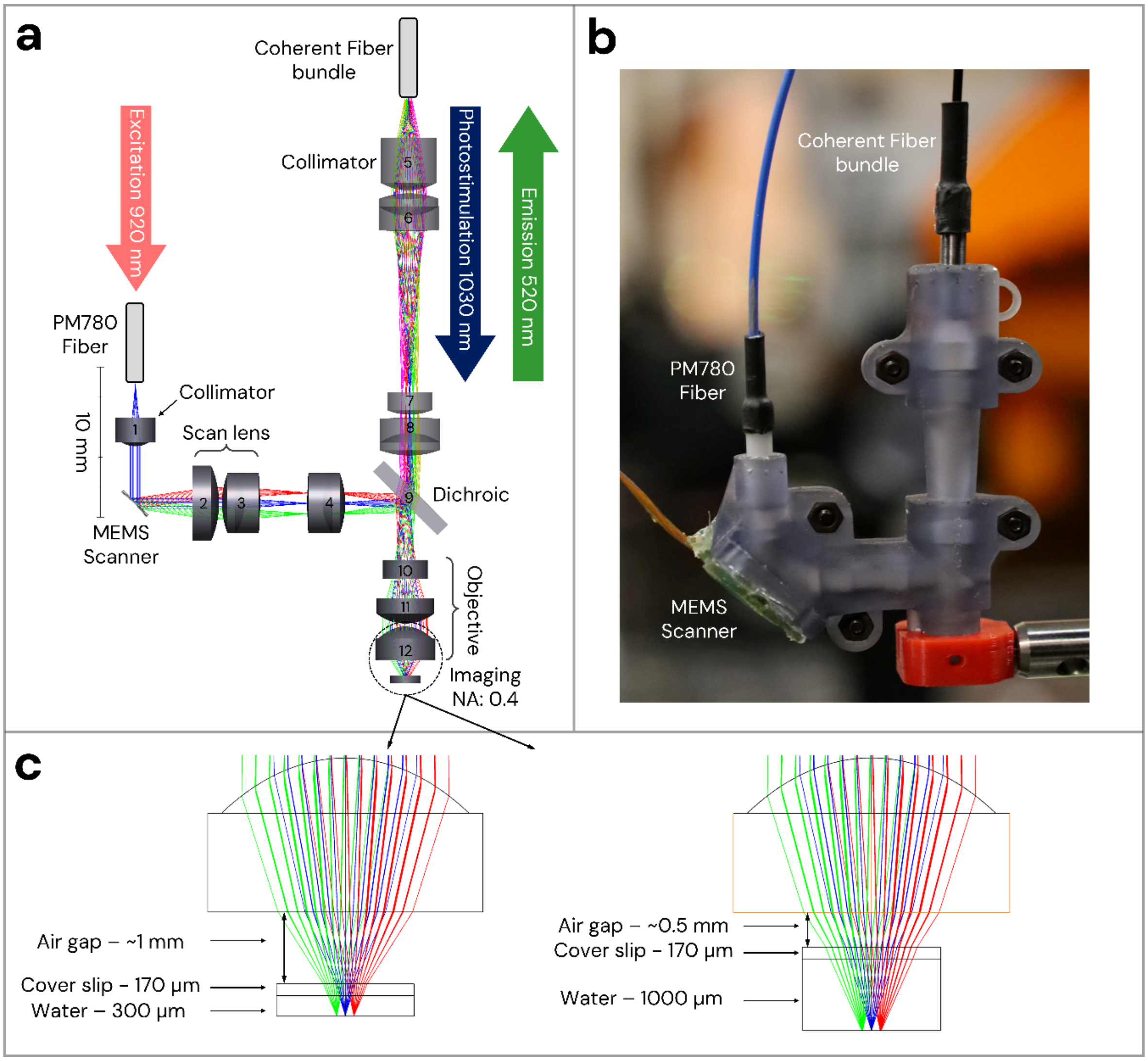


**Figure S1: Optical design, assembly and performance simulation in Zemax.** (a) Optical layout and ray tracing of the Opto2P-FCM for scanning angles of -4, 0 and 4 degrees. Red and green rays illustrate extreme angles after the MEMS scanner (±4º). Optical design consists of 920 nm excitation coming through polarization maintaining fiber (PM780), photostimulation at 1030 nm and collection coming through coherent fiber bundle (CFB). The details of the labeled optical elements are listed in Table 1. (b) Photo of assembled device in a 3D printed housing. (c) Show the minimal and maximal working distance respectively.

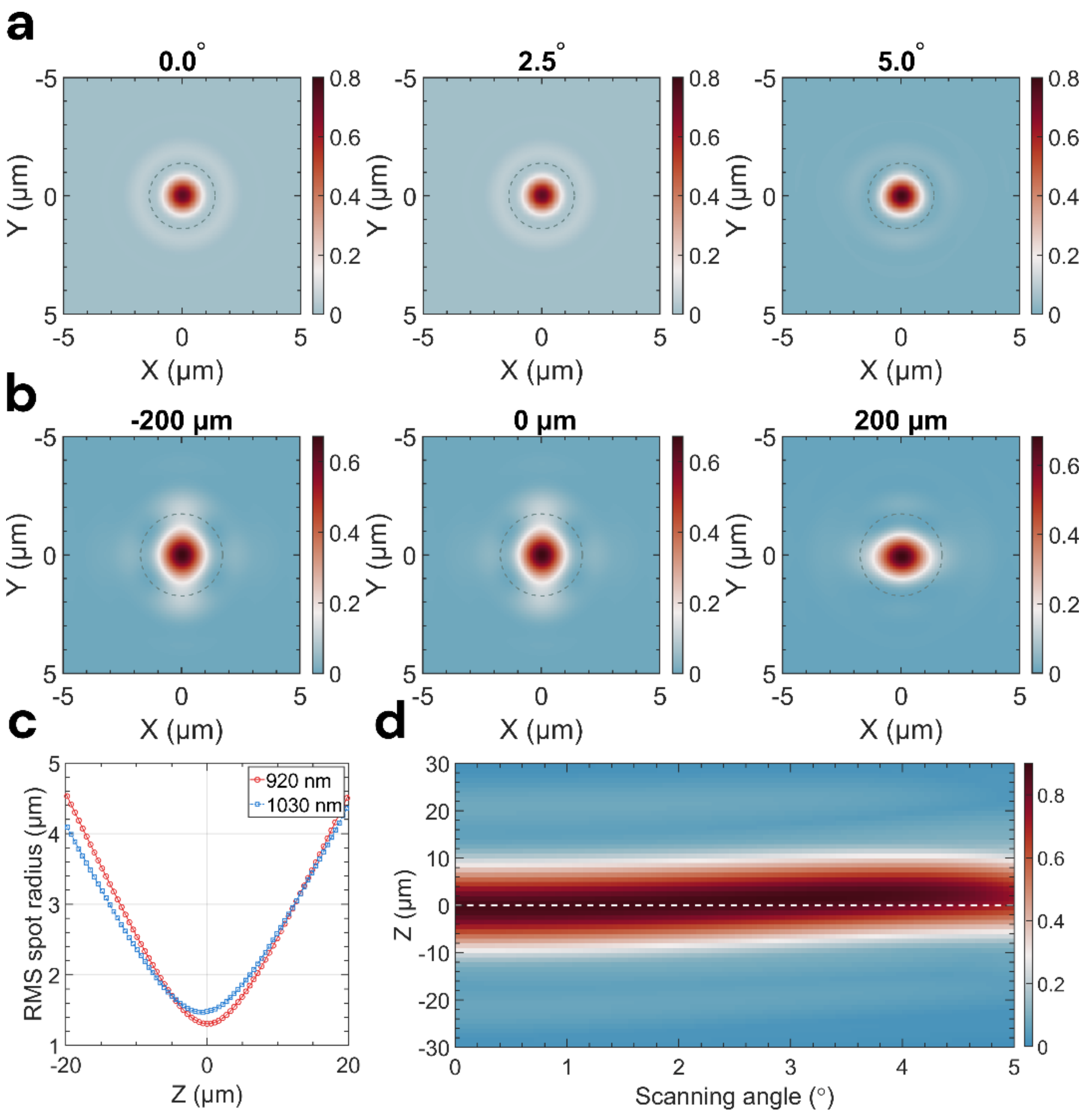


**Figure S2: Simulations of the Opto2P-FCM point-spread function.** (a) The point-spread function (PSF) of the excitation light at 920 nm for scanning angles of 0, 2.5 and 5 degrees with Strehl ratio of 0.8 and the focal spot within the Airy disk radius of 1.38 μm (dashed circle) for all scanning angles, demonstrating diffraction limit performance. (b) The PSF of 1030 nm light at the imaging plane for an input beam launched at -200, 0, and 200 μm at the entrance facet of the CFB with 0.2 NA. The dashed circle with a radius of 1.73 μm shows the Airy diffraction disk. The Strehl ratio of the photostimulation beam varies across the imaging field ranging from ~0.6 to 0.7. (c) The RMS (root-mean square) radius of the spot is plotted for excitation and photostimulation as a function of working distance with zero position on the minimum RMS spot at 920 nm. The minima of the RMS spots are offset by less than ~0.5 μm. (d) The density plot of the Strehl ratio for the excitation beam is shown in Fig. 2d as a function scanning angle of the MEMS scanner and the working distance. The system shows a 2 μm axial displacement between the 920 and 1030 nm light when scanning to 4-degree angle relative to on-axis position.

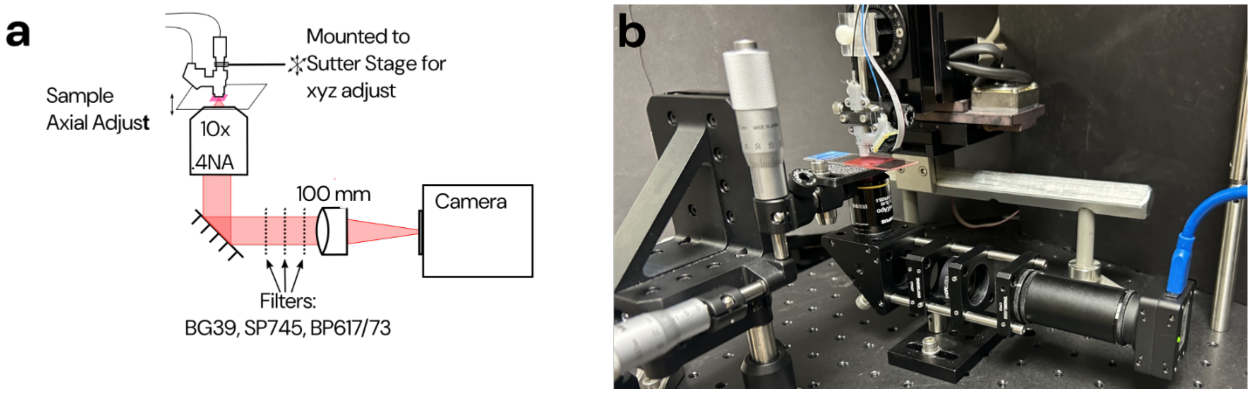


**Figure S3: Schematic of inspection scope used to perform optical characterization and spatial calibration of the imaging and photostimulation fields.** (a) Diagram of scope includes a 10x/.4 NA Olympus objective, fold mirror, filters (BG39, Short Pass 745 nm, and Band Pass 617/73 nm) for imaging fluorescence or replaced with OD filters for directly imaging the laser, a 100 mm focal length tube lens and camera (FLIR CM3-US-31S4M-CS, Edmund Optics). (b) Photograph showing inspection scope, sample holder, and Opto2P-FCM.

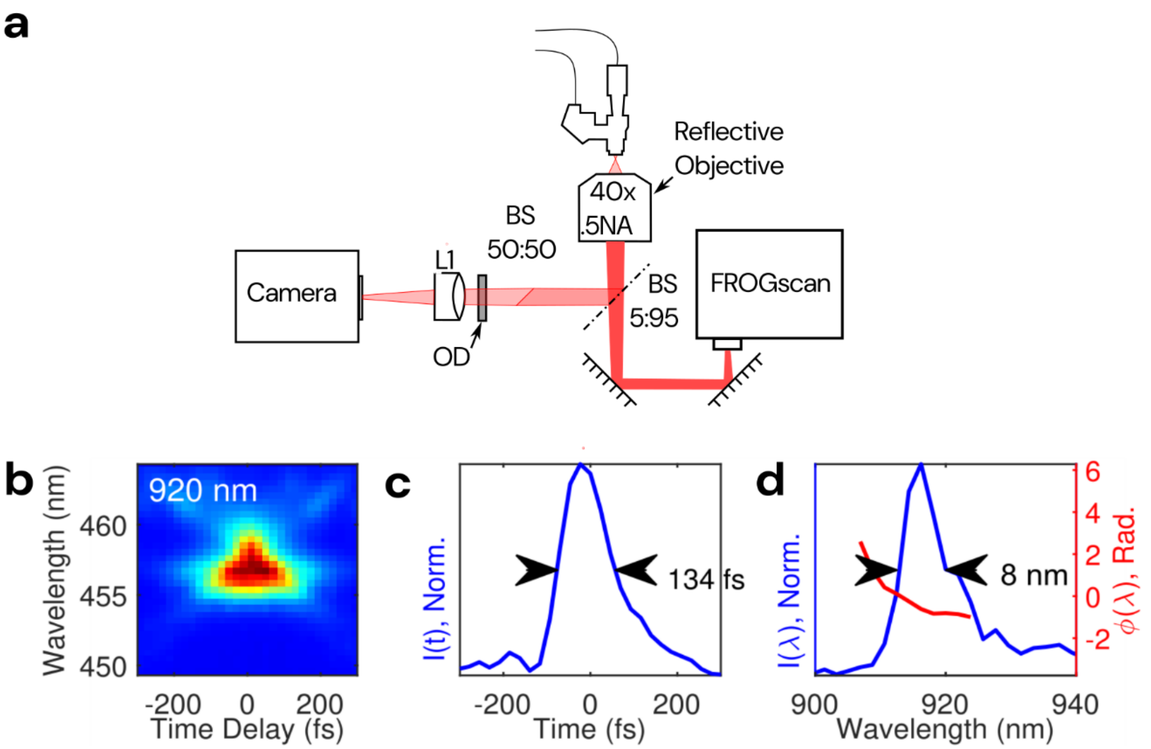


**Figure S4: Setup for pulse characterization and pulse measurements**. (a) The laser output from the Opto2P-FCM is collected by a 40x/.5 NA reflective objective (LMM40x-PO1, Thorlabs, NJ USA). Collected light is passed through a 5:95 beam splitter and relayed into a spectrally resolved autocorrelator (FROGscan, MesaPhotonics). The image of the focus is also captured on a camera using tube lens L1. (b) Frequency resolved optical gating (FROG) spectrogram of the 920 nm laser output. (c) and (d) reconstructed temporal and spectral pulse profiles with measured pulse duration of ~130 fs FWHM and bandwidth of ~8 nm FWHM.

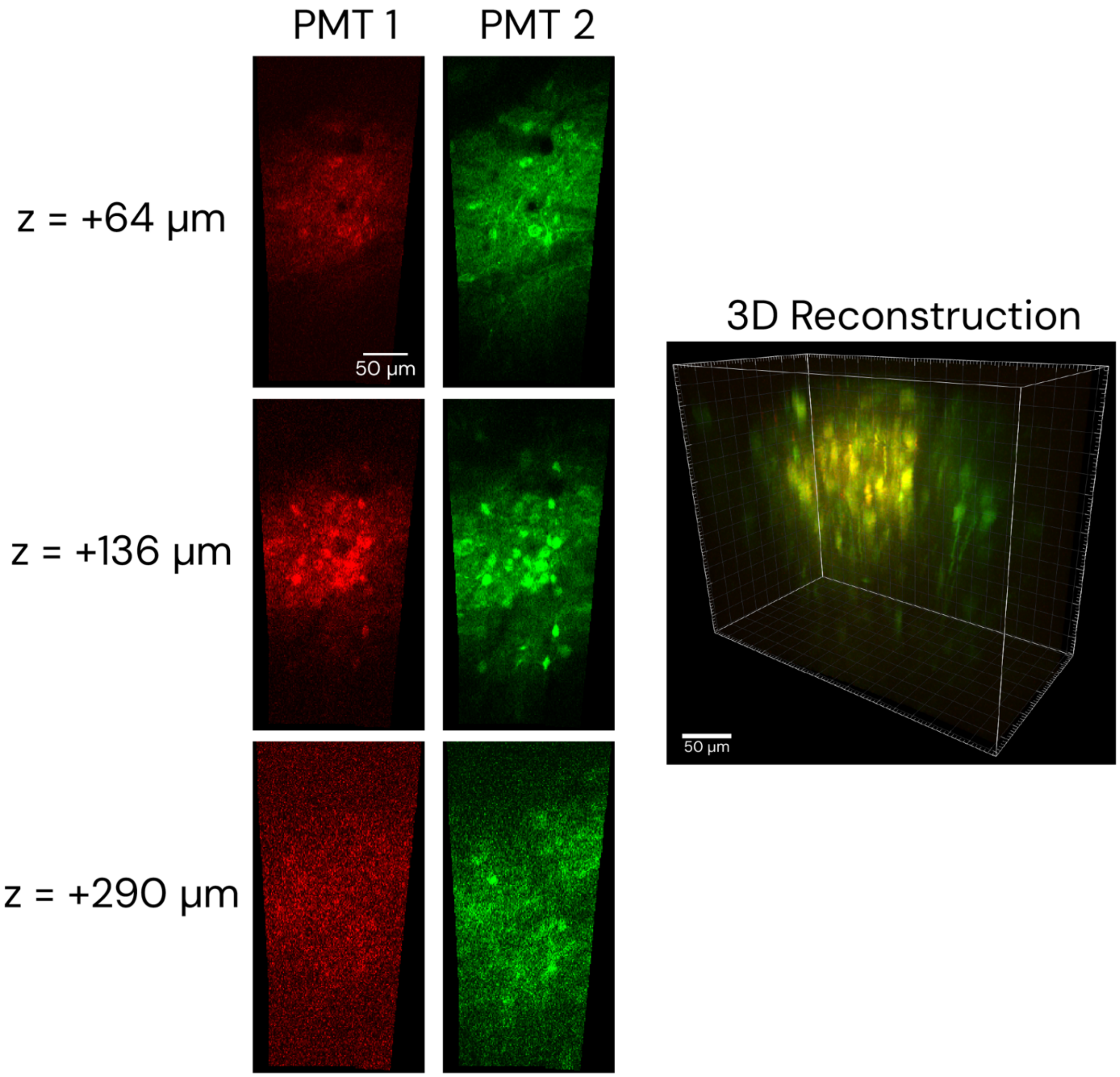


**Figure S5: Opto2P-FCM two-color imaging at depth in the visual cortex of ChRmine (Red) and jGCaMP7s (Green).** (Left) Images collected at increasing imaging depths in the red channel (PMT1) and the green channel (PMT2). Axial scanning was performed by mounting the Opto2P-FCM to a Sutter Micromanipulator stage to adjust the distance to the cranial window in a head-fixed mouse. (Right) 3D reconstruction of the imaging volume.
